# An efficient miRNA knockout approach using CRISPR-Cas9 in Xenopus

**DOI:** 10.1101/2021.08.05.454468

**Authors:** Alice M. Godden, Nicole J. Ward, Michael van der Lee, Anita Abu-Daya, Matthew Guille, Grant N. Wheeler

## Abstract

In recent years CRISPR-Cas9 knockouts (KO) have become increasingly ultilised to study gene function. MicroRNAs (miRNAs) are short non-coding RNAs, 20-22 nucleotides long, which affect gene expression through post-transcriptional repression. We previously identified miRNAs-196a and −219 as implicated in the development of *Xenopus* neural crest (NC). The NC is a multipotent stem-cell population, specified during early neurulation. Following EMT NC cells migrate to various points in the developing embryo where they give rise to a number of tissues including parts of the peripheral nervous system and craniofacial skeleton. Dysregulation of NC development results in many diseases grouped under the term neurocristopathies. As miRNAs are so small it is difficult to design CRISPR sgRNAs that reproducibly lead to a KO. We have therefore designed a novel approach using two guide RNAs to effectively ‘drop out’ a miRNA. We have knocked out miR-196a and miR-219 and compared the results to morpholino knockdowns (KD) of the same miRNAs. Validation of efficient CRISPR miRNA KO and phenotype analysis included use of whole-mount *in situ* hybridization of key NC and neural plate border markers such as *Pax3*, *Xhe2*, *Sox10* and *Snail2*, q-RT-PCR and Sanger sequencing. miRNA-219 and miR-196a KO’s both show loss of NC, altered neural plate and hatching gland phenotypes. Tadpoles show gross craniofacial and pigment phenotypes.

## INTRODUCTION

MiRNAs are short non-coding, single stranded RNAs, approximately 20-22 nucleotides in length (Alberti and Cochella, 2017; Lee et al., 1993; Shah et al., 2017). MiRNAs are initially transcribed by RNA polymerase II as a pri-miRNA stem-loop structure from the genome, which undergoes processing to form a show mature miRNA (Agarwal et al., 2015; Alberti and Cochella, 2017; Bartel, 2004; Inui et al., 2010).

MiRNAs are highly conserved between species with many orthologues discovered (Bartel, 2004). The miRNA database and repository, miRbase, currently has 2,656 mature miRNA sequences across all species. It is thought that there are >2,300 different miRNAs in humans alone (Alles et al., 2019). Recent reports suggest that 60% of all protein coding genes in mammals are regulated by one or more miRNAs (Li et al., 2018). Within the human genome it is estimated that up to 2% of genes encode for miRNAs (Miska, 2005). MiRNAs are implicated in development of various tissues in vertebrates, including chick, mouse, frog and fish (Mok et al., 2017; Ward et al., 2018); as well as in invertebrates like the worm and fruit fly (Chandra et al., 2017). Efficient methods to KO one or more miRNAs are therefore required. miRNAs can be produced from independent genes or encoded in intronic regions of the genome. They can be found in intergenic, intronic and exonic regions of the genome (Olena and Patton, 2010). The proposed CRISPR method in this paper works for intergenic and intronic miRNAs. We show this by knocking out miR-196a which is located in a *Hoxc* intron and miR-219, which is intergenic.

The NC has the potential to differentiate into many different cell types and it contributes to many tissues. The NC can migrate all over the body and become parts of the peripheral nervous system, craniofacial skeleton and pigment (Aoto et al., 2015; Cheung and Briscoe, 2003; Hatch et al., 2016). How this occurs depends on niches and environments that have the right cocktail of gene expression patterns, signals or transcription factors (Sauka-Spengler and Bronner-Fraser, 2008). MiRNAs have been suggested to play a role in NC development with Dicer KD experiments in mouse leading to NC cell death by apoptosis (Zehir et al., 2010). We have found that miR-219 and miR-196a are enriched in NC tissue, with miR-219 almost exclusively expressed in NC explants; others have also identified miR-196a implicated in eye development and NC through morpholino miRNA-KD experiments (Gessert et al., 2010; Ward et al., 2018).

CRISPR in recent years has been increasingly used in manipulating gene expression. CRISPR-Cas9 utilizes a highly specific targeted nuclease to induce genomic editing by non-homologous end joining or homology-directed repair. CRISPR is an efficient technology that can rapidly generate KO samples for analysis (Ran et al., 2013). In *X. tropicalis* Nakayama and colleagues laid the foundation and set out a simple CRISPR pipeline and use of mutations in the tyrosinase gene to generate albinism phenotypes, targeting the start codon, leading to frameshift mutation and KO (Nakayama et al., 2013). CRISPR can be used to analyse gene function, and to replicate human diseases mutations to generate mosaic targeted mutant F0’s and lines in *Xenopus* embryos (Feehan et al., 2019; Macken et al., 2021; Naert et al., 2020; Naert et al., 2017; Naert and Vleminckx, 2018; Nakayama et al., 2013).

As part of our ongoing work of looking at miRNAs in NC development we have developed a novel method to KO miRNAs quickly and efficiently in *X. tropicalis* embryos and analyse the phenotype generated transiently in the F0 population. Using this method we have begun to more clearly investigate the role of miR-196a and miR-219 in NC development.

## RESULTS AND DISCUSSION

### miRNA expression profiling

We previously identified miRNAs expressed in NC tissue through RNA-sequencing experiments on Wnt/Noggin induced animal-caps (Ward et al., 2018). Here we focus on two miRNAs identified in our earlier study; miR-196a which is located within the *Hoxc* cluster, in *HoxC9*, and miR-219 which is located intergenically. Both miRNAs have a pri-miRNA stem-loop structure that is highly conserved among the animal kingdom (Fig 1A). The initial aim was to identify when and where miR-196a and miR-219 are expressed in the developing *Xenopus* embryo. To understand when the miRNAs were expressed, q-RT-PCR was employed. Both miRNAs have a very similar profile with expression peaking initially at Nieuwkoop and Faber (NF) St.4 before dropping at gastrula stages of development and then increasing at late-gastrula and early neurula stages. Expression peaks at St.25 before dropping at tadpole stages (Fig 1B).

**Figure- 1:**
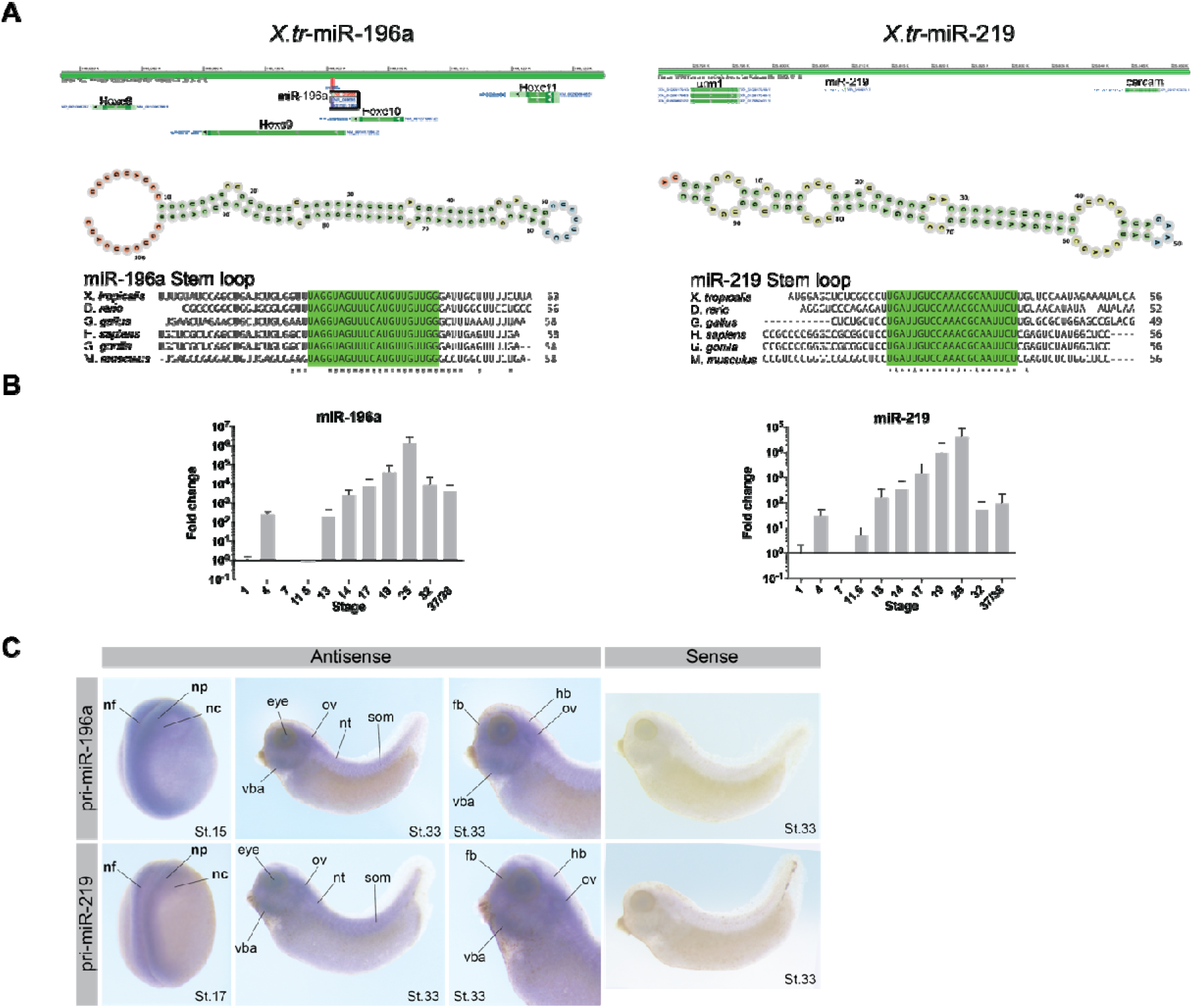
Conservation, location, spatial and temporal expression of miR-196a and miR-219 in X. *tropicalis.* *(A)* miRNA genomic locations and stem-loop structures of miRNAs. Bottom-conservation of mature miRNAs as indicated by “*”’s. miRNA stem loop structures were predicted computationally using Vienna RNA fold tool: http://rna.tbi.univie.ac.at/forna/forna.html?id=RNAfold/vCiQTz5Wd4&file=cent_probs.json (B) *X. laevis* developmental profile of miR-196a and miR-219 by qRT PCR. Fold change is represented as mean ± SD normalised to snU6 at St.1, and biological replicates with undetermined values are excluded. Ten embryos (three biological replicates) were pooled to extract total RNA for cDNA synthesis. For each biological replicate three technical replicates were conducted. (C) Spatial expression of pri-miR-196a and pri-miR-219 by whole-mount *in situ* hybridisation in X. *tropicalis* embryos at ranging stages of development. Abbreviations: nf- neural fold, np- neural plate, nc- NC, vba- ventral branchial arches, ov- otic vesicle, nt- neural tube, som- somites, fb- forebrain, hb- hindbrain.

Mature miRNAs are short 20-22 nucleotides long (Bartel, 2004). Due to this they are too short to detect with a standard *in situ* hybridisation probe (Thompson et al., 2007). We have previously used LNA modified *in situ* probes to determine miRNA expression in Xenopus embryos (Ahmed et al., 2015), however, LNA probes for miR-196a and miR-219 produced no signal (not shown). Another approach is to generate an antisense pri-miRNA *in situ* probe by PCR from genomic DNA (Walker and Harland, 2008). Using this method we looked at the expression miR-196 and miR-219. For miR-196a and miR-219 expression is very similar (Fig 1C). Expression can be seen with the antisense probes but not in sense. At neurula stages, NF St. 15 and 17, expression is seen in neural folds, neural plate and NC. At tadpole stage, NF St. 33, expression can be seen in craniofacial tissues, including NC derivatives, the otic vesicle and ventral branchial arches. Using LNA probes we also observed miR-196a and miR-219 expression in chick embryos in the neural tube, neural tissue and some expression in NC (Supplementary Fig. 1).

### Developing and employing CRISPR-Cas9

CRISPR-Cas9 approaches have been developed in many species including *Xenopus* (Nakayama et al., 2013). To KO a miRNA in *X. tropicalis*, the technical limitation of generating a viable embryo with a clean KO is that designing a sgRNA close to or in the miRNA is difficult due to their small size. In addition an insertion/deletion (INDEL) mutation could lead to generation of a novel miRNA as well as losing the orginal miRNA (Bhattacharya and Cui, 2017). Other ways to KD miRNA expression by CRISPR include targeting the DROSHA and DICER processing sites within the miRNA under study (Chang et al., 2016). Again, this is not always possible due to sgRNA design limitations due to the NGG PAM design for Cas9 used in this study (Wilson et al., 2018). Therefore, it was decided to use two sgRNAs simultaneously to “drop-out” the stem-loop of the miRNA (See Fig 2A).

**Figure- 2:**
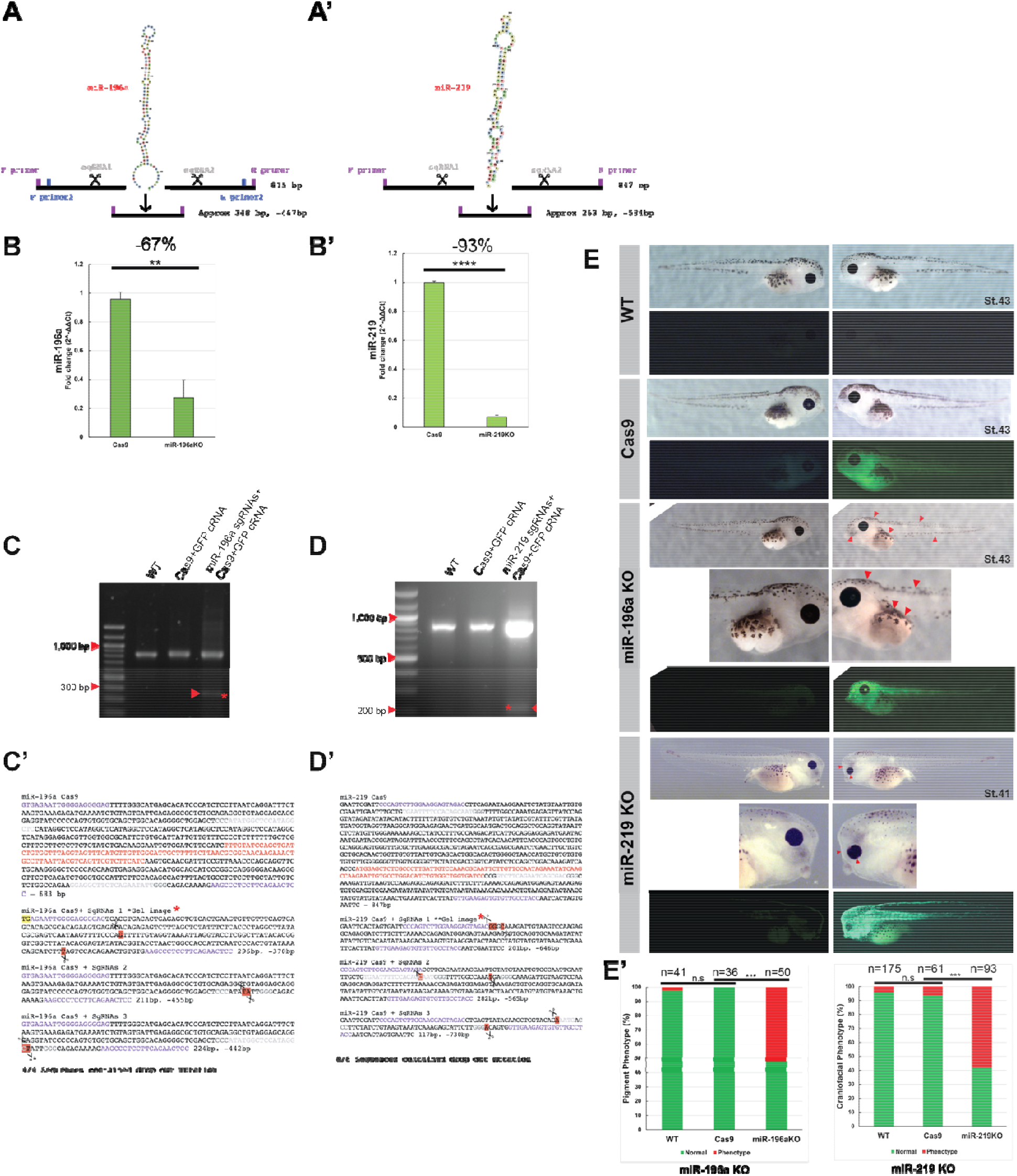
CRISPR-Cas9 approach for knocking out miRNAs in X. tropicalis and validation strategies. (A) Schematic showing the approach taken with use of two sgRNAs. (B) Q-RT-PCR validation of miR-196a KO (−67%) and miR-219 KO (−93%) reduction in miRNAs (C-D). PCR/nested PCR validation of gDNA miRNA regions from embryos injected into one cell at 2 cell stage of development, with KOs showing an extra smaller band in the fourth lane of each gel. (C’-D’) Sanger sequencing validation of miRNA KOs and CRISPR events. Cas9+GFP control samples were also harvested, genomic DNA extracted, PCR amplified and subcloned. Purple text highlights primers used for cloning, red text shows miRNA stem-loop. Yellow highlight shows a mis-match, and red highlight with scissor icons show where CRISPR events occurred, grey text shows sgRNA. WT and Cas9 sequences show miRNA WT sequence, whereas Cas9+sgRNAs show 3 repeats of mutated sequences, with significantly shorter sequences. (E) Phenotype analysis of miRNA KO embryos, representative embryos are shown. Embryos were co-injected with CRISPR-reagents a GFP capped RNA tracer into one cell at a two cell stage of development. Embryos are imaged on left and right sides. WT had no injection, Cas9 protein was co-injected with GFP cRNA and miR-KO were pairs of sgRNAs, Cas9 protein and GFP cRNA tracer. The fluorescent side of miR-196a KO embryo red arrows indicate a pigment phenotype, and for miR-219 KO, the red arrows indicate a strong craniofacial phenotype, with smaller eyes, branchial arch and flattened face features. (E’) Bar charts show count data of yes/no phenotypes for miR-196a KO (pigment loss) and miR-219 KO (craniofacial disfigurement), with chi-squared tests for statistical significance. There was a significant difference between and Cas9 and miR-196a KO groups p=2.22×10^−7^ and between Cas9 and miR-219 KO p=1.1×10^−10^. Embryo phenotypes were blind counted on three biological repeats on embryos from different frogs.

To be confident the miRNAs were knocked out, pairs of sgRNAs, Cas9 protein and GFP cRNA were co-injected into *X. tropicalis* embryos into both blastomeres at the 2-cell stage to target whole embryo (Fig 2A). Embryos expressing GFP on both sides were selected, and 5 St.14 neurulas were pooled to produce independent biological repeats and RNA was harvested to analyse miRNA expression by q-RT-PCR to evaluate sgRNA efficiency. The results showed that expression of both miRNAs was reduced in the treated samples, compared to control embryos injected with Cas9 protein and GFP cRNA tracer. miR-196a sgRNAs resulted in a 67% reduction of miR-196a expression and miR-219 sgRNAs reduced miR-219 expression by 93% as compared to control samples (Fig 2B). Differences in efficiency could be due to the nature of CRISPR (Ran et al., 2013).

Next, we identified the types of INDEL generated using the two guide-RNAs approach. Genomic DNA was extracted from individual *X. tropicalis* embryos injected with CRISPR reagents into one blastomere at 2-cell stage of development. Cas9 + tracer was used as a negative control. PCR was carried out to amplify the stem-loop of the miRNA (Fig 1A). As expected we detected wild-type (WT) miRNA in all samples. For the miR-KO samples an extra smaller band was seen on the gel (Fig 2C, D). For miR-196a the WT miRNA band is 815 bp, the sgRNAs should lead to the deletion of 467 bp and release a fragment of approximately 300 bp if a CRISPR event was successful. For miR-219 the WT miRNA should be 847 bp, the CRISPR-released fragment is expected to 260 bp or smaller, as seen in the gel (Fig 2 C & D). Amplified products were then gel extracted and sent for Sanger sequencing (Fig 2 C’ & D’). For miR-196a a nested PCR was carried out using primer set 3 (Table 2). Stem-loops for each miRNA are shown by red text, and as expected the “drop-out” bands do not contain the miRNA, and confirm the successful CRISPR deletion of miRNA stem-loops (Fig 2 C’ & D’). The WT and Cas9 control group bands were also extracted and sent for sequencing these all showed wild-type miRNA sequences. This to our knowledge, is the first time this approach with two sgRNAs has been used in *Xenopus* embryos, though Kretov et al (2020) do report a similar approach in Zebrafish (Kretov et al., 2020).

Some embryos were left to develop into tadpoles for phenotype analysis. Embryos were targeted with CRISPR reagents on one side only for comparison with the non-injected side as an internal control. WT and Cas9 embryos look morphologically normal on both sides. However, miR-196a tadpoles on the “crispant” (CRISPR-mutated) side show pigment phenotypes, with a reduction in pigment seen along the cranial, dorsal and medial abdominal regions, as indicated by red arrows (Fig. 2E). MiR-219 tadpoles show gross craniofacial impairments, as shown by the red arrows (Fig. 2E). Blind counts of the phenotypes showed that over 50% of embryos carried the respective pigment and craniofacial phenotypes (Fig 2E’). These phenotypes suggest a possible role for miR-196a and miR-219 in NC development (Collazo et al., 1993; Lukoseviciute et al., 2018; Petratou et al., 2021; Scerbo and Monsoro-Burq, 2020; Spokony et al., 2002).

### Exploring miRNA phenotypes

To verify if our miRNAs were implicated in the development of NC, we determined the expression of key markers Sox10, Snail2 for NC, Pax3 for neural plate and Xhe2 for hatching gland (Fig 3). In addition, we compared the efficiency of our CRISPR KO approach versus morpholino mediated KD of the miRNAs. Control and optimization experiments for miRNA-morpholino KD can be seen in Suppl. Fig. 2.

**Figure- 3:**
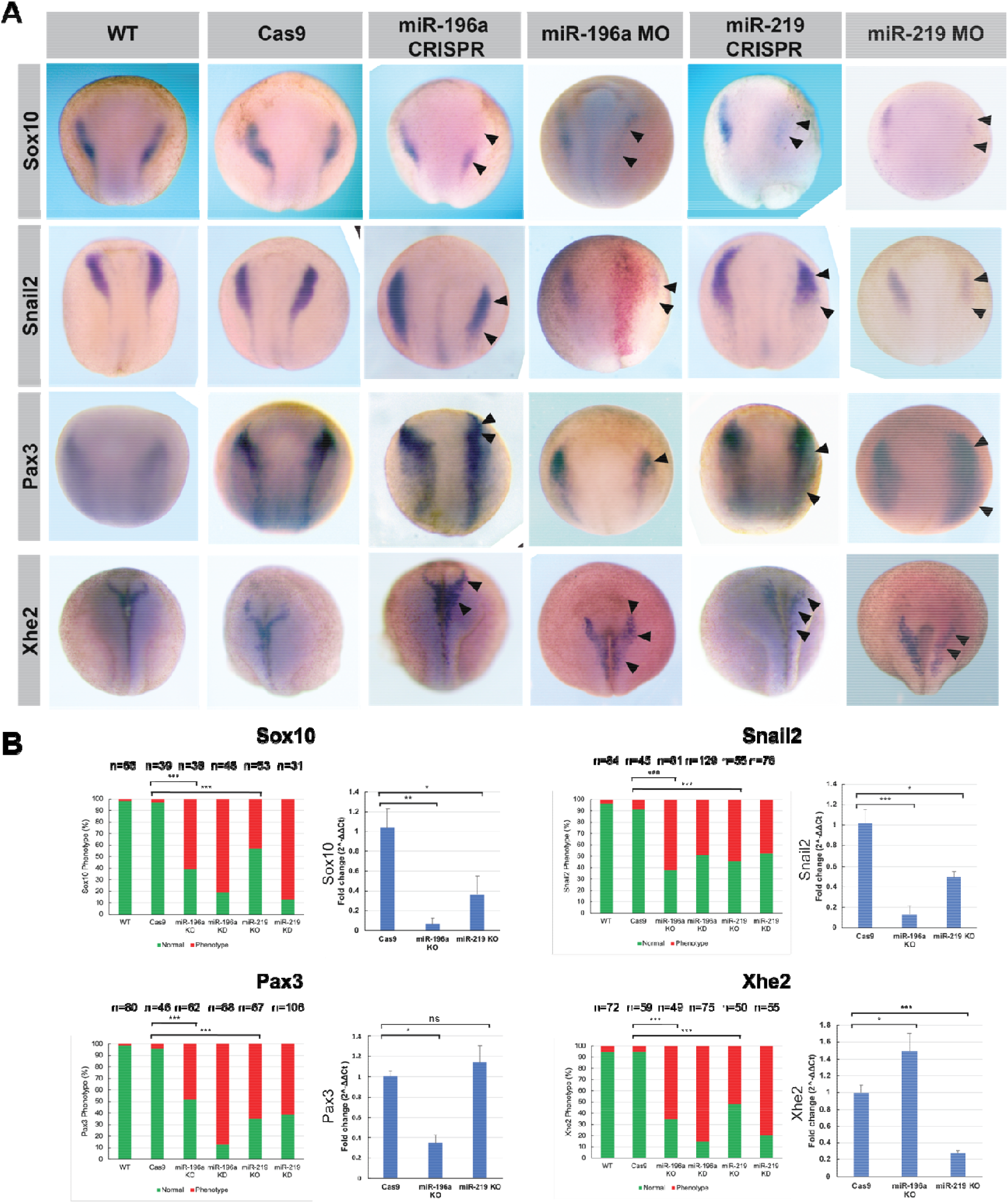
Analysis of key NC, neural plate and hatching gland markers after miRNA KO and KD. (A) Whole mount *in situ* hybridisation profiles on neurula stage *Xenopus* embryos of Sox10, Snail2, Pax3 and Xhe2. CRISPR-Cas9 was carried out in *X. tropicalis* embryos with GFP cRNA as a tracer and morpholino-KD was carried out in *X. laevis* embryos with lacZ cRNA as a tracer. Embryos for whole mount *in situ* hybridisation were injected with tracer at 4-cell stage of development into the right dorsal blastomere. Overall phenotypes show a reduction of NC and altered neural plate and hatching gland profiles. (B) Phenotype analysis for individual markers. Bar charts show count data of yes/no phenotype. Chi-squared statistical tests were carried out on three biological repeats of whole mount *in situ* hybridisation on embryos from different frogs. q-RT-PCR was carried out for mRNA expression analysis. Normalised to U6 expression, RNA was pooled from 5 individual neurula embryos for one biological sample, q-RT-PCR was carried out with biological and technical triplicates. q-RT-PCR data supports phenotypes shown in (A). Abbreviations for phenotype and q-RT-PCR bar charts in (B): miR-KO refers to CRISPR miRNA KO and miR- KD refers to MO KD of miRNA. Phenotypes for miRNA KD/KO: Sox10 phenotype is a reduction in expression, Snail2 is a reduction/shift in profile, Pax3 phenotype is a shift/reduced profile for miR-196a and an expansion for miR-219 experiments, finally, Xhe2 is an increased profile for miR-196a and a reduced profile for miR-219 experiments respectively. Statistical significance: Sox10 Cas9 vs miR-196a KO p= 4.02 × 10^−8^, Cas9 vs miR-219 KO p=1.04 × 10^−5^, Snail2 Cas9 vs miR-196a KO p= 6.15 × 10^−9^, Cas9 vs miR-219 KO p=4.07 × 10^−7^, Pax3 Cas9 vs miR-196a KO p= 7.19 × 10^−7^, Cas9 vs miR-219 KO p=2.29 × 10^−8^, Xhe2 Cas9 vs miR-196a KO p= 7.19 × 10^−7^, Cas9 vs miR-219 KO p=2.29 × 10^−8^.

For miR-196a KO and KD led to distinct reduction in Sox10 expression. For Snail2 miR-196 KO and KD led to a reduction in expression showing a shift in expression. For Pax3 expression a slight reduction and shift in profile was observed and finally Xhe2 expression was expanded following miR-196a KO and KD (Fig. 3A-B). For miR-219 KO and KD, again Sox10 expression was markedly decreased. Like with miR-196 KD, Snail2 expression following miR-219 KD was reduced, but miR-219 KO showed more of a shift in profile with slight reduction. Following miR-219 KO and KD Pax3 expression was greatly increased and expanded (Fig 3A-B), section data showed miR-219 KD led to Pax3 expansion over the superficial ectoderm but not following miR-196a KD (Suppl. Fig. 3). Phenotypes shown by *in situ* hybridisation were more prevalent for miRNA-KD than KO, this could be due to potential mosaicism of CRISPR events seen in F0 embryos thus leading to variable miRNA levels between embryos (Naert et al., 2020).

q-RT-PCR was conducted to quantify the phenotypic change in expression of the above markers across the whole embryo (Fig 3B). q-RT-PCR was carried out on crispant samples, as described previously, injecting CRISPR reagents into one blastomere at the 1 cell stage. For mRNA expression, all the q-RT-PCR data was in agreement with the in situ data except for the miR-219 KO on Pax3, which was shown to be increased in expression though this was not significant (Fig 3B). Reasons for this could be that the whole embryo was used for RNA extraction. It may be that overall there isn’t a huge change in Pax3 expression, and that it is shifted to superficial ectoderm (Suppl. Fig. 3). The data indicate that miR-196a and miR-219 could be involved in early regulation of NC development.

The incidence rate of phenotypes can be seen in Fig 3B with the observed phenotypes clearly occurring in the experimental groups. Broadly miRNA KD and KO phenotype incidence were similar between morpholino and CRISPR, although miR-196a morpholino was notably higher rate of phenotype incidence for Sox10 and Pax3. q-RT-PCR profiles match the *in situ* data. With NC markers showing significant decreases in expression, more notably for miR-196a. Pax3 expansion for miR-219 KO was not statistically significant, but miR-196a KO led to significant reduction in expression. Xhe2 showed significant expansion for miR-196a KO and significant decrease in miR-219 KO. These results show that miRNAs are likely to be implicated in the development of the *Xenopus* NC and can be analysed through use of CRISPR to KO miRNAs.

The loss of Sox10 expression shown in Fig. 3A for miR-196a and miR-219 KD and KO supports the phenotypes shown in the tadpoles in Fig. 2E. The tadpole phenotypes for miR-196a show loss of pigment, and for miR-219 show craniofacial abnormalities, which is significant as Sox10 is involved in trunk NC to produce pigmentation (Aoki et al., 2003) and is disrupted in neurocristopathies affecting cranial NC (Devotta et al., 2016).

Snail2 expression is downregulated following loss of miRNA (Fig. 3A). This could mean that miRNA KO and KD is leading to a loss of NC differentiation, and may help explain the craniofacial phenotypes seen in miR-219 KO tadpoles, (Fig. 2E), (Li et al., 2019). The loss of pigment phenotype seen following miR-196a KO (Fig. 2E), are typical of problems in trunk NC development (Abu-Elmagd et al., 2006). This is also supported by Snail2^−/−^ mice that show patchy pigmentation phenotypes (Shi et al., 2011).

The loss of Pax3 seen in Fig. 3A following loss of miR-196a also supports the pigment phenotype seen in Fig. 2E, as Pax3 is essential for the development of pigment (Kubic et al., 2008). Waardenburg syndrome type 1 and 3 are caused by Pax3 gene mutations, and type 2 and 4 are affected by Sox10 and MITF levels and mutations. Pax3 and Sox10 regulate expression of MITF and thus melanocyte development (Bondurand et al., 2000). As expected our results follow this by showing reduced Pax3 and Sox10 expression following miR-196 loss (Fig. 3A) which led to pigment loss in tadpoles (Fig. 2E). The expansion of Pax3 following loss of miR-219 (Fig 3A), could be affecting NC specification leading to the loss of NC induction expression of Snail2 and Sox10 seen in Fig. 3A(McKeown et al., 2005; Monsoro-Burq et al., 2005).

The expansion of Pax3 observed in superficial ectoderm (Suppl Fig. 3) following miR-219 KD could also be indicative of increased neural pluripotency (Chalmers et al., 2002). *Xenopus* neural plate border region can give rise to placodal ectoderm, hatching gland and NC. The hatching gland marker Xhe2 demarcates this. Xhe2 expression is affected by Pax3 expression (Hong and Saint-Jeannet, 2007; Hong and Saint-Jeannet, 2014), miR-196a and miR-219 KO and KD experiments show altered Pax3 expression states, therefore as expected Xhe2 expxression was also affected. In contrast to results seen in (Hong and Saint-Jeannet, 2014), Pax3 expansion does not lead to expanded Xhe2 in our work, this could be due to the other signals that mediate Xhe2 like Zic1. Further work is required to investigate more markers across neural plate and neural deriviatives.

## CONCLUSIONS

One drawback of CRISPR gene-editing is finding sgRNAs that are effective especially when studying miRNAs. As mentioned, a challenge with CRISPR experiments is that with a short target sequence the number of sgRNAs that can be designed is limited (Najah et al., 2019). With the advent of more Cas9 nucleases with broader PAM recognition sequences more designs could be generated (Kim et al., 2020). Furthermore new sgRNA design tools are making sgRNA design easier and more robust (Hsu et al., 2021).However, using sgRNAs flanking the miRNA stem-loop expands the potential for identifying and generating optimal sgRNAs. Using a pair of sgRNAs leads to a complete loss of the miRNA in the majority of embryos. The method described is a quick and efficient way to KO specific miRNAs in independent genes or within introns. With the generation of lines of frogs, time would be saved from laborious injections of morpholinos and controls thus more ambitious and technically demanding experiments would be more realistic.

This work shows that miRNAs miR-196a and miR-219 are expressed in NC and neural tissue. Phenotype analysis shows that the miRNAs are important for NC and hatching gland development. This body of work goes some way to establishing protocols and controls for CRISPR experiments for knocking out miRNAs in embryo development and puts forward CRISPR as not only a tool to rival use of morpholinos in embryo research but also to potentially replace in certain instances.

## MATERIALS AND METHODS

### Xenopus husbandry

All experiments were carried out in accordance with relevant laws and institutional guidelines at the University of East Anglia, with full ethical review and approval, compliant to UK Home Office regulations. Embryos were generated as described in (Harrison et al., 2004; Williams et al., 2017). *X. tropicalis* embryos obtained by priming females up to 72 hrs before use with 10 ui chorulon and induced on the day of use with 200 ui. Eggs were collected manually and fertilised in vitro. Embryos were de-jellied in 2% L-cysteine, incubated at 23°C and microinjected in 3% Ficoll into 1 cell at the 2-4 cell stage in the animal pole. Embryos were left to develop at 23°C. Embryo staging is according to Nieuwkoop and Faber (NF) normal table of Xenopus development (Nieuwkoop, 1967). GFP/LacZ capped RNA for injections was prepared using the SP6 mMESSAGE mMACHINE kit, 5 ng was injected per embryo.

### CRISPR-Cas9

SgRNAs were designed using CRISPRScan (https://www.crisprscan.org/), (Moreno-Mateos et al., 2015). miRNA sequences were attained from miRbase (http://www.mirbase.org/); under accession numbers: Xtr-miR-219 MI0004873, Xtr-miR-196a MI0004942. SgRNAs were designed up and downstream of the miRNA stem-loop. miRNA stem loop structures were predicted computationally using Vienna RNA fold tool (http://rna.tbi.univie.ac.at/forna/forna.html?id=RNAfold/vCiQTz5Wd4&file=cent_probs.json).

#### sgRNAs

**Table 1.**
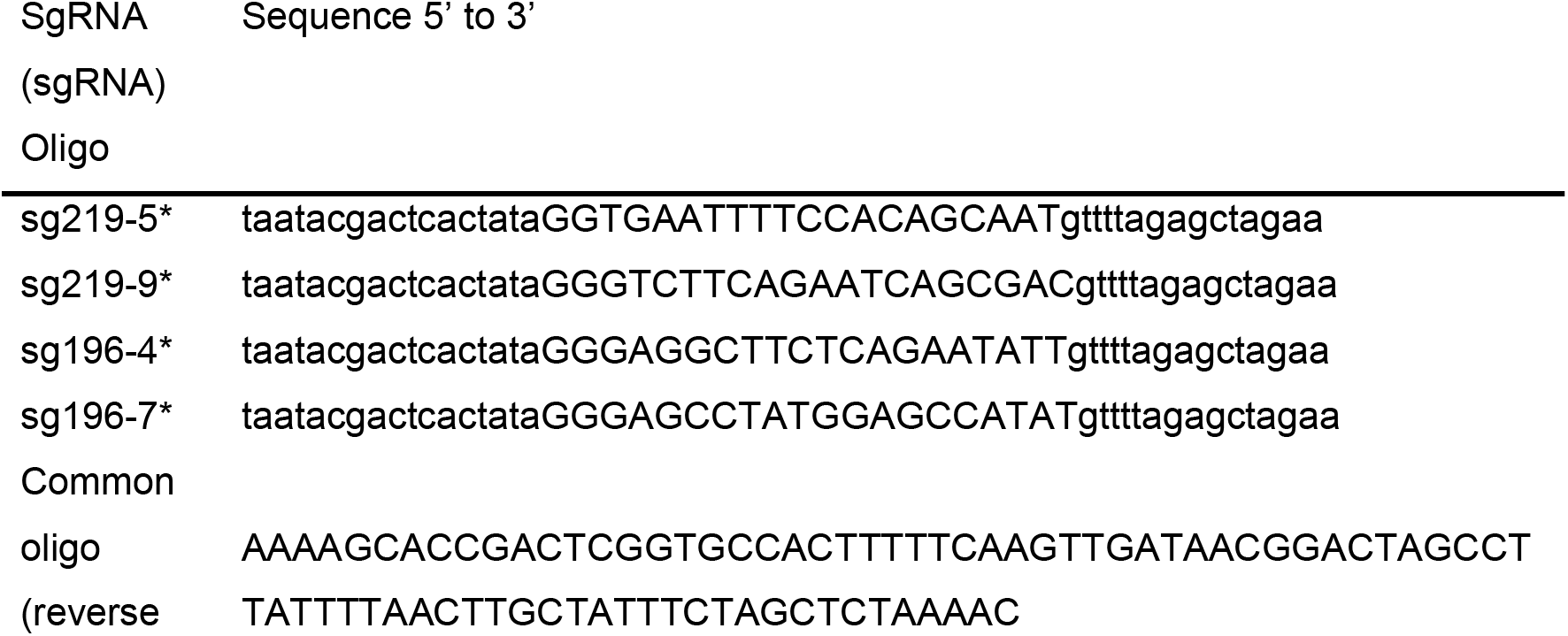
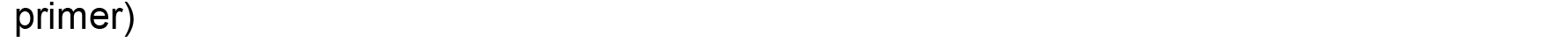
SgRNA sequences used. Common oligo taken from (Nakayama et al., 2013).

**Table 2.**
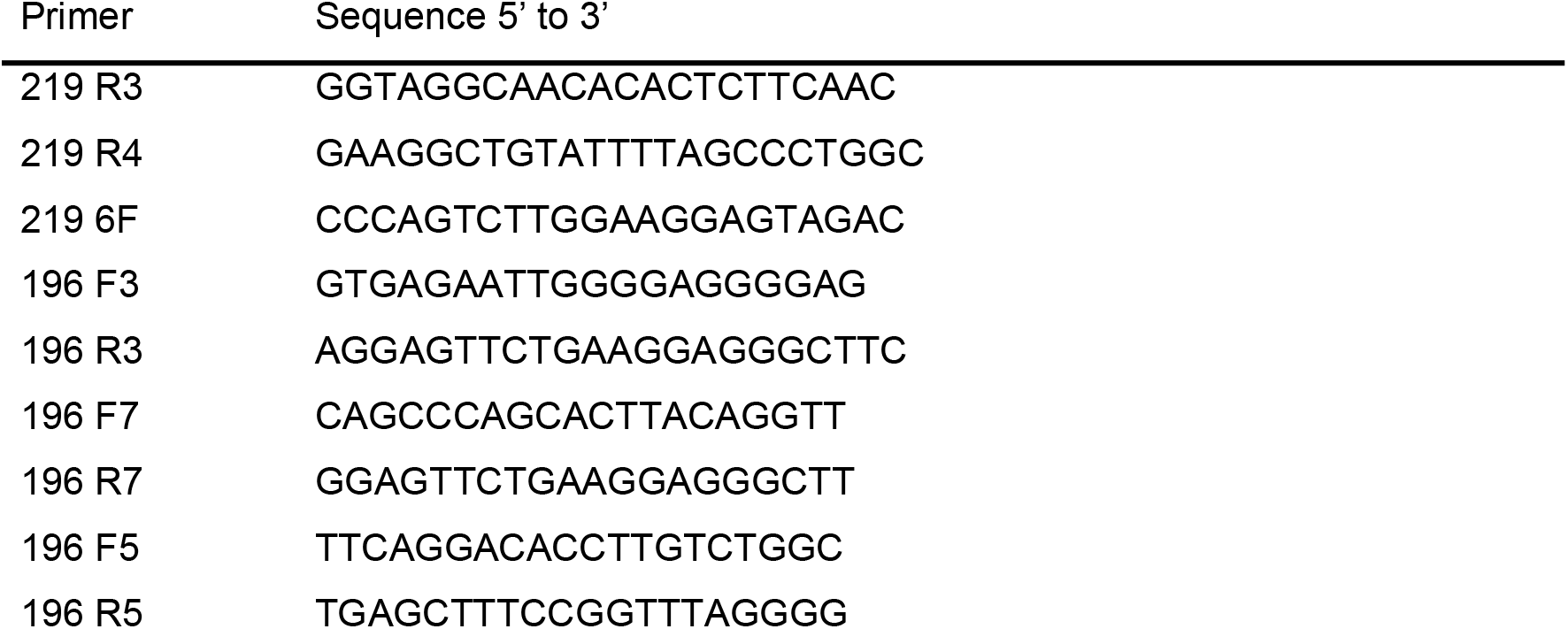
Table of primers and sequences. Primers used for sequencing and PCR of genomic DNA. Designed using Primer3.

### Embryo injection

Embryos were injected using a 10 nL calibrated needle. For *X. laevis* 10 nL injections, for *X. tropicalis* 4.2 nL injections were used. Cas9 protein (New England Biolabs, #M0646M, EnGen Cas9 NLS 20 μM) and 300 pg of sgRNAs along with 5 ng of GFP cRNA were co-injected into the *X. tropicalis* embryos simultaneously at 2-4 cell stage of development. For q-RT-PCR both sides of embryo were targeted, for gene expression and morphological analysis phenotypes 1 side of the embryo was targeted, with embryos injected at 4 cell stage into one dorsal blastomere for whole-mount in situ experiments and morphological analysis.

### CRISPR Validation

Embryos were left to develop until tadpole stages and underwent phenotype scoring. Embryos were then frozen on dry ice before genomic DNA extraction. Genomic DNA was isolated using PureLink Genomic DNA Mini Kit, K1820-00 (Invitrogen, California, USA), according to manufacturers guidelines and then quantified using a Nanodrop 1000. Genotyping PCRs were conducted and products underwent gel extraction before subcloning and sanger sequencing.

#### Morpholinos

Morpholino dose was optimized to 60 ng for miRNAs; morpholino and lacZ capped RNA tracer were injected at 4 cell stage of embryo development into the right dorsal blastomere.

**Table 3.**
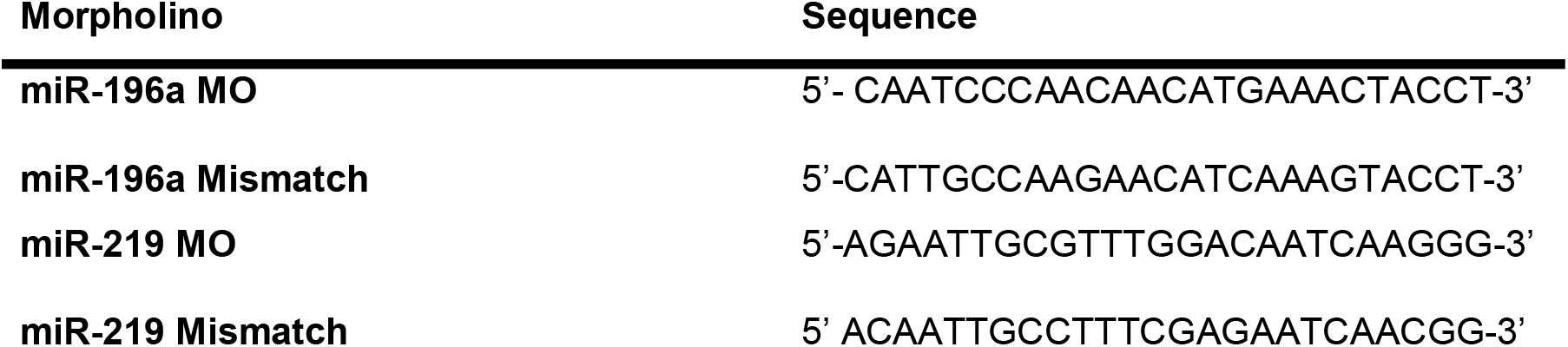
Injected morpholino sequence data

### Phenotype Statistical analysis

Chi-squared test for association was used to test phenotype yes or no categories for morpholino injected embryos to see if there was a relationship between two categorical values. Excel was used to collate and tabulate data. IBM SPSS v25 to carry out chi-squared test. Statistical significance is reported as; p<0.05 = *, p<0.01 = **, p<0.001 = ***.

### q-RT-PCR

Embryos were frozen on dry ice before RNA extraction. For miRNA and mRNA quantification total RNA was extracted from five St.14 X. *tropicalis* embryos, embryos were homogenised with a micropestle and RNA was extracted according to manufacturers guidance, Quick-RNA Mini prep plus kit (Zymo, Cat no. R1058). Samples were eluted in 25 μL of nuclease free water; RNA concentration and purity quantified on a Nanodrop 1000 and 1 μL was checked on a 2% agarose gel. All q-RT-PCR’s were performed with triplicate biological and technical repeats.

### cDNA synthesis

To produce cDNA for q-RT-PCR, miRCURY LNA RT kit (Qiagen, Cat No./ID: 339340). 50 ng of RNA was used and kit according to manufacturers instructions. cDNA was produced on a thermocycler with the following programme: 42 °C for 60 min and 95 °C for 5 min. cDNA was diluted 1:40 for q-RT-PCR. CDNA can be stored at −20°C. To produce cDNA for mRNA q-RT-PCR the following recipe was used: 500 ng of total RNA was added in 9 uL of nuclease free water, plus 2 uL of random primers (Promega, C1181). This was then incubated at 70 °C for 10 mins. A mastermix was prepared as follows per sample: 4 uL of 5X buffer, 2 uL of DTT, 1 uL of dNTPs, 1 uL of Superscript II (Invitrogen, 18064014), 1 uL of nuclease free water or RNasin (Promega, N2611). qRT-PCR reactions were set up in 10 μL volume containing 4 μL cDNA, 1 μL primer (10 μM for standard oligo primers), and 5 μL SybrGreen (Applied Biosystems 4309155).

Primers for q-RT-PCR were found in the literature and some were designed using primer blast (https://www.ncbi.nlm.nih.gov/tools/primer-blast/), (Ye et al., 2012). Primers were designed to generate 100 bp products with a melting temperature of between 59-62 °C. Primers used are listed in Table 4.

**Table 4.**
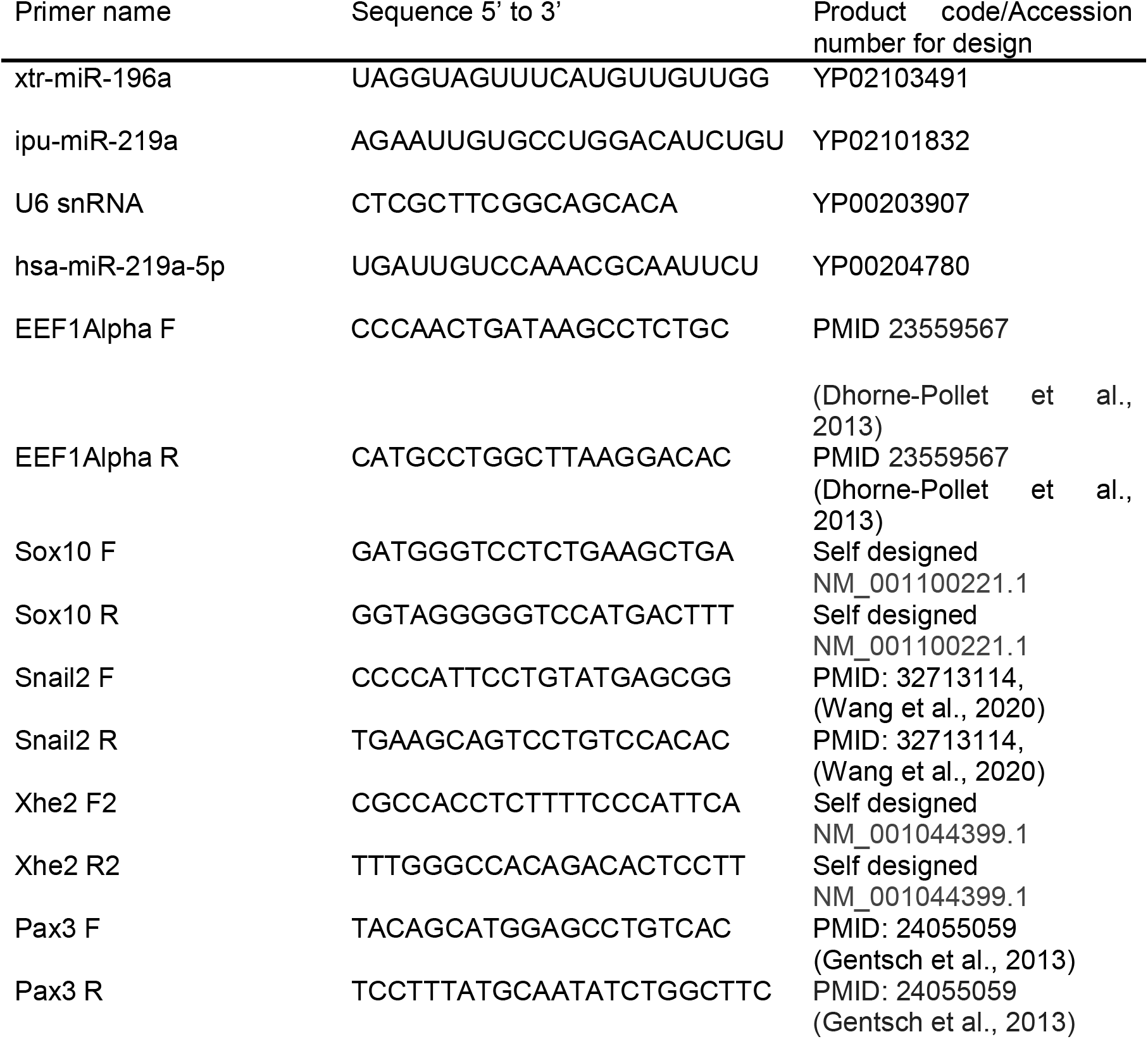
q-RT PCR Primers used for Xenopus Tropicalis embryos. miRCURY LNA miRNA PCR primers, Qiagen. mRNA primers were ordered as standard oligos.

#### Whole-mount in situ hybridisation & Riboprobe synthesis

Whole-mount in situ hybridisation with LNA probes was carried out according to (Ahmed et al., 2015; Sweetman et al., 2006). Other in situs and probe synthesis were carried out according to (Harrison et al., 2004; Monsoro-Burq, 2007; Sive et al., 2007). Plasmids with sources are listed in Table 5.

**Table 5.** Riboprobe synthesis and capped RNA plasmids information

| **Clone name** | **Antibiotic resistance** | **Backbone** | **Antisense RE** | **Antisense Polymerase** | **Source** |
| --- | --- | --- | --- | --- | --- |
| Pax3 | Ampicillin | pBSK | BglII | SP6 | Michael G. Sargent |
| Sox10 | Ampicillin | pBSK | EcoRI | T3 | JP. Saint-Jeannet |
| Snail2 | Ampicillin | pCS107 | EcoRI/BamHI | T7 | EXRC |
| Xhe2 | Ampicillin | pBSK | XbaI | T7 | AH.Monsoro-Burq |
| GFP2 | Ampicillin | pCS | NotI (sense) | SP6 (sense) | Dr. Maggie Walmesly |
| LacZ | Ampicillin | pCS | NotI (sense) | SP6 (sense) | Dr. Maggie Walmesly |

## Acknowledgements

We would like to acknowledge to ongoing technical support from the team at the European Xenopus Resource Centre at the University of Portsmouth. We also acknowledge the support from: Dr. Timothy Grocott for chick embryo expertise, Paige Paddy and Dr. Tracey Swingler for q-RT-PCR advice and Dr. Benjamin Rix for useful CRISPR discussions. We would like to thank Prof Monsoro-Burq (Orsay), Prof Saint-Jeannet (New York), Dr. Sargent (MRC) and Dr Walmsely (Nottingham) for sending plasmids. We would like to acknowledge Prof Münsterberg and Prof Moxon for help with experimental design and the Wheeler/ Münsterberg/ Grocott labs for all their support.

## Abbreviations

INDEL: Insertion, deletion (mutation)
miRNAs: miRNAs
KD: Knockdown
KO: Knockout
NC: Neural crest
NF: Nieuwkoop and Faber
sgRNA: single guide RNA

## SUPPLEMENTAL FIGURES

**Supplementary Figure- 1:**
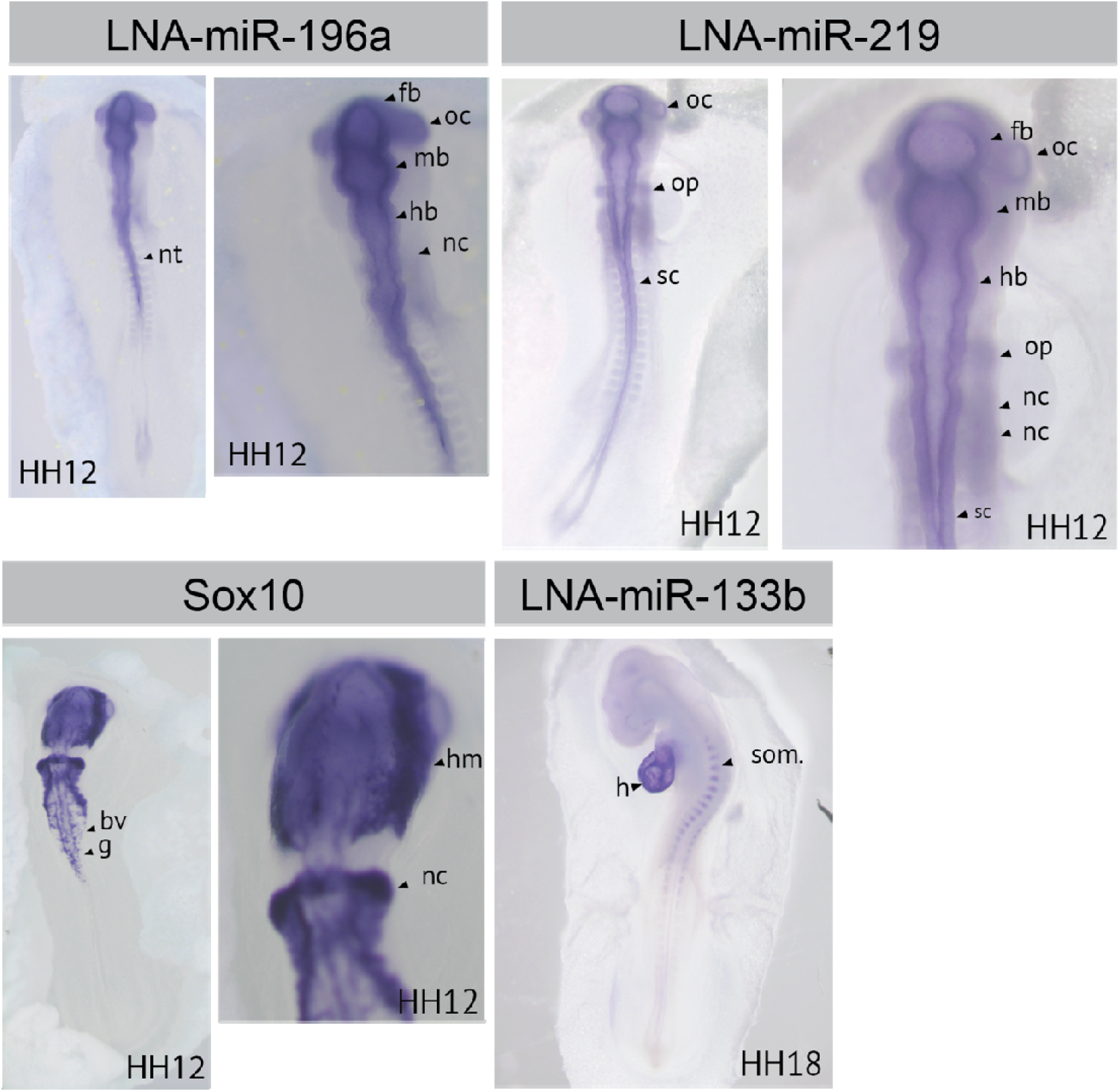
Expression profile of: Sox10, miR-196a, miR-219 and miR-133b in chick embryos. Embryos staged according to Hamilton & Hamburger. Sox10 expression can clearly be seen at HH12. Sox10 is expressed in the blood vessels, ganglia, head mesenchyme but also clearly in the NC, as indicated by black arrows. miRNA in situs were carried out with LNA probes. LNA-miR-133b clearly shows expression in the heart and somites at HH18. LNA-miR-196a and LNA-miR-219 both show expression in NC tissues, and neural tissues such as brain structures. Abbreviations: bv- blood vessels, fb- forebrain, h- heart, g- ganglia, hb- hindbrain, hm- head mesenchyme, mb- midbrain, nc- NC, ne- neural, np- neural plate, nt- neural tube, oc- optic cup, op- otic placode, sc- spinal cord, som- somites.

**Supplementary Figure- 2:**
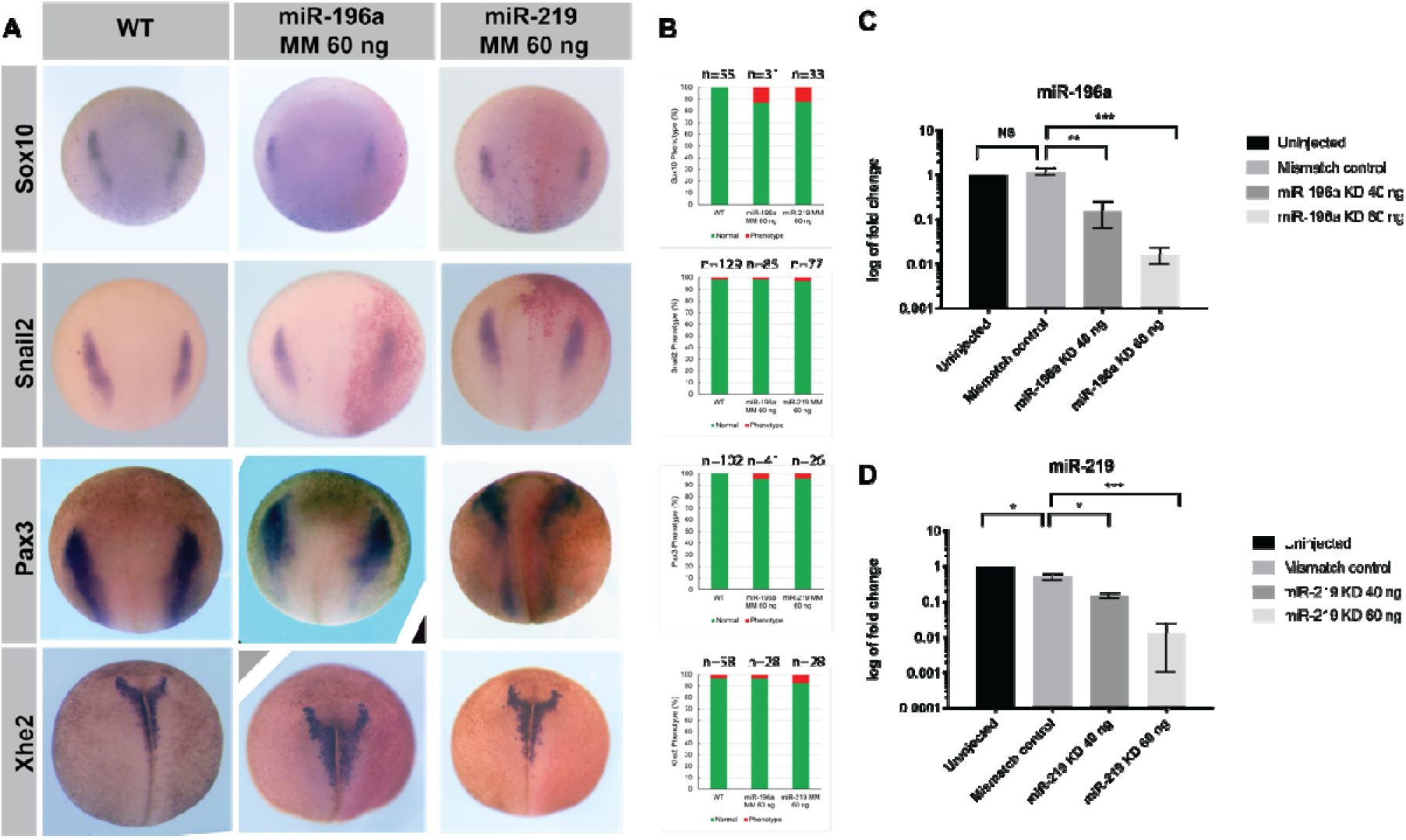
Control and validation experiments for morpholino-mediated miRNA KD. (A) Control experiments with morpholinos, showing WT, control morpholino “MM”-mismatch, for miR-196a and miR-219. X. *laevis* embryos were injected at 4-cell stage into the right dorsal blastomere with LacZ cRNA tracer. (B) Shows phenotype count data, assessed as in Fig.3. (C) q-RT-PCR validation of morpholino dose-response to show morpholino specificity. One-way Anova with post-hoc Tukey tests show statistical significance, with p=<0.05 for “*” as the threshold, p=<0.01 “**” and p=<0.001 for “***”, and ns= not significant.

**Supplementary Figure- 3:**
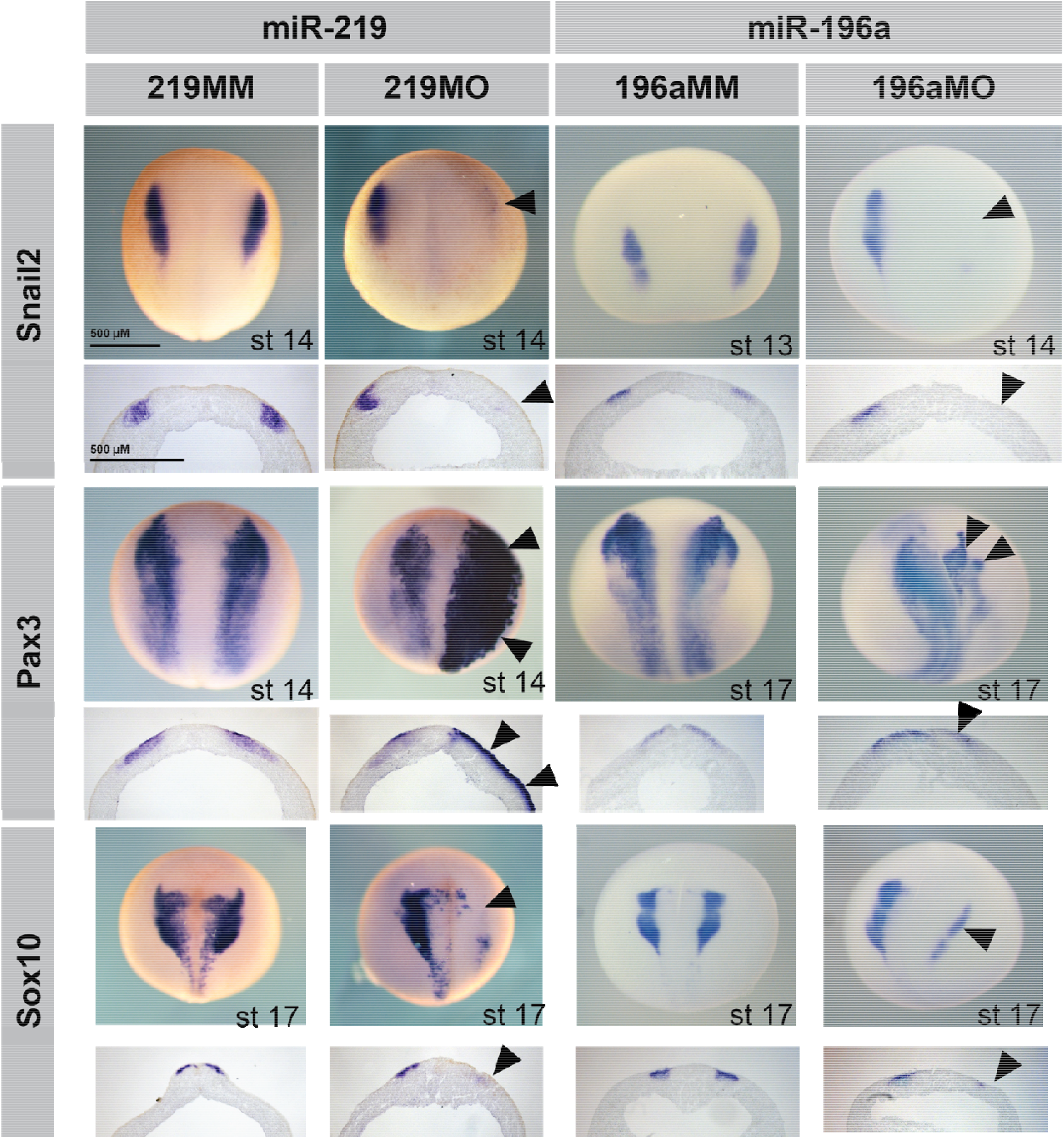
Morpholino-mediated KD of miRNAs and expression of key markers. 219MM= miR-219 mismatch-morpholino 60 ng, 219MO= miR-219 morpholino, 196MM= miR-196a mismatch-morpholino 60 ng, 196MO= miR-196a morpholino. GFP cRNA tracer was used, and positive embryos were selected for sectioning. NC markers Snail2 and Sox10 show clear reduction in expression whereas neural plate marker Pax3 shows marked expansion in superficial ectoderm region for miR-219 KD, whereas miR-196a KD shows altered expression.

## REFERENCES

Abu-Elmagd, M., Garcia-Morales, C. and Wheeler, G. N. (2006). Frizzled7 mediates canonical Wnt signaling in neural crest induction. Dev Biol 298, 285–298.

Agarwal, V., Bell, G. W., Nam, J. W. and Bartel, D. P. (2015). Predicting effective microRNA target sites in mammalian mRNAs. Elife 4.

Ahmed, A., Ward, N. J., Moxon, S., Lopez-Gomollon, S., Viaut, C., Tomlinson, M. L., Patrushev, I., Gilchrist, M. J., Dalmay, T., Dotlic, D., et al. (2015). A Database of microRNA Expression Patterns in Xenopus laevis. PLoS One 10, e0138313.

Alberti, C. and Cochella, L. (2017). A framework for understanding the roles of miRNAs in animal development. Development 144, 2548–2559.

Alles, J., Fehlmann, T., Fischer, U., Backes, C., Galata, V., Minet, M., Hart, M., Abu-Halima, M., Grasser, F. A., Lenhof, H. P., et al. (2019). An estimate of the total number of true human miRNAs. Nucleic Acids Res 47, 3353–3364.

Aoki, Y., Saint-Germain, N., Gyda, M., Magner-Fink, E., Lee, Y. H., Credidio, C. and Saint-Jeannet, J. P. (2003). Sox10 regulates the development of neural crest-derived melanocytes in Xenopus. Dev Biol 259, 19–33.

Aoto, K., Sandell, L. L., Butler Tjaden, N. E., Yuen, K. C., Watt, K. E., Black, B. L., Durnin, M. and Trainor, P. A. (2015). Mef2c-F10N enhancer driven beta-galactosidase (LacZ) and Cre recombinase mice facilitate analyses of gene function and lineage fate in neural crest cells. Dev Biol 402, 3–16.

Bartel, D. P. (2004). MicroRNAs: genomics, biogenesis, mechanism, and function. Cell 116, 281–297.

Bhattacharya, A. and Cui, Y. (2017). Systematic Prediction of the Impacts of Mutations in MicroRNA Seed Sequences. J Integr Bioinform 14.

Bondurand, N., Pingault, V., Goerich, D. E., Lemort, N., Sock, E., Le Caignec, C., Wegner, M. and Goossens, M. (2000). Interaction among SOX10, PAX3 and MITF, three genes altered in Waardenburg syndrome. Hum Mol Genet 9, 1907–1917.

Chalmers, A. D., Welchman, D. and Papalopulu, N. (2002). Intrinsic differences between the superficial and deep layers of the Xenopus ectoderm control primary neuronal differentiation. Dev Cell 2, 171–182.

Chandra, S., Vimal, D., Sharma, D., Rai, V., Gupta, S. C. and Chowdhuri, D. K. (2017). Role of miRNAs in development and disease: Lessons learnt from small organisms. Life Sci 185, 8–14.

Chang, H., Yi, B., Ma, R., Zhang, X., Zhao, H. and Xi, Y. (2016). CRISPR/cas9, a novel genomic tool to knock down microRNA in vitro and in vivo. Sci Rep 6, 22312.

Cheung, M. and Briscoe, J. (2003). Neural crest development is regulated by the transcription factor Sox9. Development 130, 5681–5693.

Collazo, A., Bronner-Fraser, M. and Fraser, S. E. (1993). Vital dye labelling of Xenopus laevis trunk neural crest reveals multipotency and novel pathways of migration. Development 118, 363–376.

Devotta, A., Juraver-Geslin, H., Gonzalez, J. A., Hong, C. S. and Saint-Jeannet, J. P. (2016). Sf3b4-depleted Xenopus embryos: A model to study the pathogenesis of craniofacial defects in Nager syndrome. Dev Biol 415, 371–382.

Dhorne-Pollet, S., Thelie, A. and Pollet, N. (2013). Validation of novel reference genes for RT-qPCR studies of gene expression in Xenopus tropicalis during embryonic and post-embryonic development. Dev Dyn 242, 709–717.

Feehan, J. M., Stanar, P., Tam, B. M., Chiu, C. and Moritz, O. L. (2019). Generation and Analysis of Xenopus laevis Models of Retinal Degeneration Using CRISPR/Cas9. Methods Mol Biol 1834, 193–207.

Gentsch, G. E., Owens, N. D., Martin, S. R., Piccinelli, P., Faial, T., Trotter, M. W., Gilchrist, M. J. and Smith, J. C. (2013). In vivo T-box transcription factor profiling reveals joint regulation of embryonic neuromesodermal bipotency. Cell Rep 4, 1185–1196.

Gessert, S., Bugner, V., Tecza, A., Pinker, M. and Kuhl, M. (2010). FMR1/FXR1 and the miRNA pathway are required for eye and neural crest development. Dev Biol 341, 222–235.

Harrison, M., Abu-Elmagd, M., Grocott, T., Yates, C., Gavrilovic, J. and Wheeler, G. N. (2004). Matrix metalloproteinase genes in Xenopus development. Dev Dyn 231, 214–220.

Hatch, V. L., Marin-Barba, M., Moxon, S., Ford, C. T., Ward, N. J., Tomlinson, M. L., Desanlis, I., Hendry, A. E., Hontelez, S., van Kruijsbergen, I., et al. (2016). The positive transcriptional elongation factor (P-TEFb) is required for neural crest specification. Dev Biol 416, 361–372.

Hong, C. S. and Saint-Jeannet, J. P. (2007). The activity of Pax3 and Zic1 regulates three distinct cell fates at the neural plate border. Mol Biol Cell 18, 2192–2202.

Hong, C. S. and Saint-Jeannet, J. P. (2014). Xhe2 is a member of the astacin family of metalloproteases that promotes Xenopus hatching. Genesis 52, 946–951.

Hsu, J. Y., Grunewald, J., Szalay, R., Shih, J., Anzalone, A. V., Lam, K. C., Shen, M. W., Petri, K., Liu, D. R., Keith Joung, J., et al. (2021). PrimeDesign software for rapid and simplified design of prime editing guide RNAs. Nat Commun 12, 1034.

Inui, M., Martello, G. and Piccolo, S. (2010). MicroRNA control of signal transduction. Nat Rev Mol Cell Biol 11, 252–263.

Kim, H. K., Lee, S., Kim, Y., Park, J., Min, S., Choi, J. W., Huang, T. P., Yoon, S., Liu, D. R. and Kim, H. H. (2020). High-throughput analysis of the activities of xCas9, SpCas9-NG and SpCas9 at matched and mismatched target sequences in human cells. Nat Biomed Eng 4, 111–124.

Kretov, D. A., Walawalkar, I. A., Mora-Martin, A., Shafik, A. M., Moxon, S. and Cifuentes, D. (2020). Ago2-Dependent Processing Allows miR-451 to Evade the Global MicroRNA Turnover Elicited during Erythropoiesis. Mol Cell 78, 317–328 e316.

Kubic, J. D., Young, K. P., Plummer, R. S., Ludvik, A. E. and Lang, D. (2008). Pigmentation PAX-ways: the role of Pax3 in melanogenesis, melanocyte stem cell maintenance, and disease. Pigment Cell Melanoma Res 21, 627–645.

Lee, R. C., Feinbaum, R. L. and Ambros, V. (1993). The C. elegans heterochronic gene lin-4 encodes small RNAs with antisense complementarity to lin-14. Cell 75, 843–854.

Li, J., Perfetto, M., Materna, C., Li, R., Thi Tran, H., Vleminckx, K., Duncan, M. K. and Wei, S. (2019). A new transgenic reporter line reveals Wnt-dependent Snai2 re-expression and cranial neural crest differentiation in Xenopus. Sci Rep 9, 11191.

Li, Z., Xu, R. and Li, N. (2018). MicroRNAs from plants to animals, do they define a new messenger for communication? Nutr Metab (Lond) 15, 68.

Lukoseviciute, M., Gavriouchkina, D., Williams, R. M., Hochgreb-Hagele, T., Senanayake, U., Chong-Morrison, V., Thongjuea, S., Repapi, E., Mead, A. and Sauka-Spengler, T. (2018). From Pioneer to Repressor: Bimodal foxd3 Activity Dynamically Remodels Neural Crest Regulatory Landscape In Vivo. Dev Cell 47, 608–628 e606.

Macken, W. L., Godwin, A., Wheway, G., Stals, K., Nazlamova, L., Ellard, S., Alfares, A., Aloraini, T., AlSubaie, L., Alfadhel, M., et al. (2021). Biallelic variants in COPB1 cause a novel, severe intellectual disability syndrome with cataracts and variable microcephaly. Genome Med 13, 34.

McKeown, S. J., Lee, V. M., Bronner-Fraser, M., Newgreen, D. F. and Farlie, P. G. (2005). Sox10 overexpression induces neural crest-like cells from all dorsoventral levels of the neural tube but inhibits differentiation. Dev Dyn 233, 430–444.

Miska, E. A. (2005). How microRNAs control cell division, differentiation and death. Curr Opin Genet Dev 15, 563–568.

Mok, G. F., Lozano-Velasco, E. and Munsterberg, A. (2017). microRNAs in skeletal muscle development. Semin Cell Dev Biol 72, 67–76.

Monsoro-Burq, A. H. (2007). A rapid protocol for whole-mount in situ hybridization on Xenopus embryos. CSH Protoc 2007, pdb prot4809.

Monsoro-Burq, A. H., Wang, E. and Harland, R. (2005). Msx1 and Pax3 cooperate to mediate FGF8 and WNT signals during Xenopus neural crest induction. Dev Cell 8, 167–178.

Moreno-Mateos, M. A., Vejnar, C. E., Beaudoin, J. D., Fernandez, J. P., Mis, E. K., Khokha, M. K. and Giraldez, A. J. (2015). CRISPRscan: designing highly efficient sgRNAs for CRISPR-Cas9 targeting in vivo. Nat Methods 12, 982–988.

Naert, T., Tulkens, D., Edwards, N. A., Carron, M., Shaidani, N. I., Wlizla, M., Boel, A., Demuynck, S., Horb, M. E., Coucke, P., et al. (2020). Maximizing CRISPR/Cas9 phenotype penetrance applying predictive modeling of editing outcomes in Xenopus and zebrafish embryos. Sci Rep 10, 14662.

Naert, T., Van Nieuwenhuysen, T. and Vleminckx, K. (2017). TALENs and CRISPR/Cas9 fuel genetically engineered clinically relevant Xenopus tropicalis tumor models. Genesis 55.

Naert, T. and Vleminckx, K. (2018). Methods for CRISPR/Cas9 Xenopus tropicalis Tissue-Specific Multiplex Genome Engineering. Methods Mol Biol 1865, 33–54.

Najah, S., Saulnier, C., Pernodet, J. L. and Bury-Mone, S. (2019). Design of a generic CRISPR-Cas9 approach using the same sgRNA to perform gene editing at distinct loci. BMC Biotechnol 19, 18.

Nakayama, T., Fish, M. B., Fisher, M., Oomen-Hajagos, J., Thomsen, G. H. and Grainger, R. (2013). Simple and efficient CRISPR/Cas9-mediated targeted mutagenesis in Xenopus tropicalis. Genesis 51, 835–843.

Nieuwkoop, P., Faber, J., (1967). Normal Table of Xenopus Laevis (Daudin): A Systematical and Chronological Survey of the Development from the Fertilized Egg Till the End of Metamorphosis. New York: Garland Publishing.

Olena, A. F. and Patton, J. G. (2010). Genomic organization of microRNAs. J Cell Physiol 222, 540–545.

Petratou, K., Spencer, S. A., Kelsh, R. N. and Lister, J. A. (2021). The MITF paralog tfec is required in neural crest development for fate specification of the iridophore lineage from a multipotent pigment cell progenitor. PLoS One 16, e0244794.

Ran, F. A., Hsu, P. D., Wright, J., Agarwala, V., Scott, D. A. and Zhang, F. (2013). Genome engineering using the CRISPR-Cas9 system. Nat Protoc 8, 2281–2308.

Sauka-Spengler, T. and Bronner-Fraser, M. (2008). A gene regulatory network orchestrates neural crest formation. Nat Rev Mol Cell Biol 9, 557–568.

Scerbo, P. and Monsoro-Burq, A. H. (2020). The vertebrate-specific VENTX/NANOG gene empowers neural crest with ectomesenchyme potential. Science Advances 6, In Press.

Shah, V. V., Soibam, B., Ritter, R. A., Benham, A., Oomen, J. and Sater, A. K. (2017). MicroRNAs and ectodermal specification I. Identification of miRs and miR-targeted mRNAs in early anterior neural and epidermal ectoderm. Dev Biol 426, 200–210.

Shi, J., Severson, C., Yang, J., Wedlich, D. and Klymkowsky, M. W. (2011). Snail2 controls mesodermal BMP/Wnt induction of neural crest. Development 138, 3135–3145.

Sive, H. L., Grainger, R. M. and Harland, R. M. (2007). Synthesis and purification of digoxigenin-labeled RNA probes for in situ hybridization. CSH Protoc 2007, pdb prot4778.

Spokony, R. F., Aoki, Y., Saint-Germain, N., Magner-Fink, E. and Saint-Jeannet, J. P. (2002). The transcription factor Sox9 is required for cranial neural crest development in Xenopus. Development 129, 421–432.

Sweetman, D., Rathjen, T., Jefferson, M., Wheeler, G., Smith, T. G., Wheeler, G. N., Munsterberg, A. and Dalmay, T. (2006). FGF-4 signaling is involved in mir-206 expression in developing somites of chicken embryos. Dev Dyn 235, 2185–2191.

Thompson, R. C., Deo, M. and Turner, D. L. (2007). Analysis of microRNA expression by in situ hybridization with RNA oligonucleotide probes. Methods 43, 153–161.

Walker, J. C. and Harland, R. M. (2008). Expression of microRNAs during embryonic development of Xenopus tropicalis. Gene Expr Patterns 8, 452–456.

Wang, C., Qi, X., Zhou, X., Sun, J., Cai, D., Lu, G., Chen, X., Jiang, Z., Yao, Y. G., Chan, W. Y., et al. (2020). RNA-Seq analysis on ets1 mutant embryos of Xenopus tropicalis identifies microseminoprotein beta gene 3 as an essential regulator of neural crest migration. FASEB J 34, 12726–12738.

Ward, N. J., Green, D., Higgins, J., Dalmay, T., Munsterberg, A., Moxon, S. and Wheeler, G. (2018). microRNAs associated with early neural crest development in Xenopus laevis. BMC Genomics 19, 59.

Williams, R. M., Winkfein, R. J., Ginger, R. S., Green, M. R., Schnetkamp, P. P. and Wheeler, G. N. (2017). A functional approach to understanding the role of NCKX5 in Xenopus pigmentation. PLoS One 12, e0180465.

Wilson, L. O. W., O’Brien, A. R. and Bauer, D. C. (2018). The Current State and Future of CRISPR-Cas9 gRNA Design Tools. Front Pharmacol 9, 749.

Ye, J., Coulouris, G., Zaretskaya, I., Cutcutache, I., Rozen, S. and Madden, T. L. (2012). Primer-BLAST: a tool to design target-specific primers for polymerase chain reaction. BMC Bioinformatics 13, 134.

Zehir, A., Hua, L. L., Maska, E. L., Morikawa, Y. and Cserjesi, P. (2010). Dicer is required for survival of differentiating neural crest cells. Dev Biol 340, 459–467.

